# Computing temporal sequences associated with dynamic patterns on the C. elegans connectome

**DOI:** 10.1101/2020.05.08.085191

**Authors:** Vivek Kurien George, Francesca Puppo, Gabriel A. Silva

## Abstract

Understanding how the structural connectivity of a network constrains the dynamics it is able to support is a very active and open area of research. We simulated the plausible dynamics resulting from the known C. elegans connectome using a recent model and theoretical analysis that computes the dynamics of neurobiological networks by focusing on how local interactions among connected neurons give rise to the global dynamics in an emergent way, independent of the biophysical or molecular details of the cells themselves. We studied the dynamics which resulted from stimulating a chemosensory neuron (ASEL) in a known feeding circuit, both in isolation and embedded in the full connectome. We show that contralateral motor neuron activations in ventral (VB) and dorsal (DB) classes of motor neurons emerged from the simulations, which are qualitatively similar to rhythmic motor neuron firing pattern associated with locomotion of the worm. One interpretation of these results is that there is an inherent - and we propose - purposeful structural wiring to the C. elegans connectome that has evolved to serve specific behavioral functions. To study network signaling pathways responsible for the dynamics we developed an analytic framework that constructs Temporal Sequences (TSeq), time-ordered walks of signals on graphs. We found that only 5% of TSeq are preserved between the isolated feeding network relative to its embedded counterpart. The remaining 95% of signaling pathways computed in the isolated network are not present in the embedded network. This suggests a cautionary note for computational studies of isolated neurobiological circuits and networks.

## 1 Introduction

A variety of network analyses methods are widely used to study complex systems [49, 64, 21, 7, 9, 10, 59]. But understanding how the structural connectivity of a network constrains the dynamics it is able to support is still an active and open area of research. This is particularly the case for spatial-temporal networks, where much of the literature and methods are still in early stages and mostly descriptive, providing few tools for predictive modeling. As a result, the application of this emerging body of work towards the treatment and analyses of biological neural networks that are inherently spatial-temporal remains quite limited. But the theoretical and practical insights that such an approach would offer neurobiology are potentially very significant. One such application is the analyses of the dynamics that is supported by a given structural connectome. This remains at the forefront of neuroscience research [6, 14]. In this work we introduce a novel set of methods and analyses for this task, and apply it to the dynamics of the Caenorhabditis elegans worm connectome as an example.

We first simulated the plausible dynamics resulting from the known C. elegans connectome [25, 31, 18] using a recent model and theoretical study we published that computes the dynamics of neurobiological networks by focusing on how local interactions among connected neurons give rise to the global dynamics in an emergent way, independent of the biophysical or molecular details of the cells themselves [58]. This framework was derived from a theoretical abstraction of the canonical principles of spatial and temporal summation in biological neurons. By analyzing the arrival times of incident signals into nodes and how signals compete to activate downstream nodes, we were able to prove a number of properties associated with the dynamics of the network. Our model takes into account how the timing of the arrival of different signals on target neurons influences how they interact to activate the nodes they connect into, given that the internal (refractory) state of the target neuron they are competing for may or may not allow the cell to be activated. Following a series of simulations, the main contribution of this work was the development of a set of novel methods and tools that allowed us to extract the spatial-temporal patterns and activation sequences of nodes that make up the dynamics of the network. In essence, this allows us to understand how local rules drive global change in order to analyze the casual interactions and patterns of node activations that produced the computed dynamics of the network.

The C. elegans connectome consists of the anatomical edges (links) between its 302 neurons, and was fully mapped in 1986 [67]. Although the organism’s connectome has been known for decades, how neurons functionally interact in the context of the entire network and how the resultant dynamics regulates participating neurons is not fully understood [6, 5, 39]. Recent research efforts have moved past analyzing the network’s structure [64, 62], towards analyzing its dynamics in various ways [35, 11, 55, 48, 61, 70, 36]. Here we asked how does the concurrent activity of independent neuronal elements ultimately give rise to a rich behavioral repertoire of the C. elegans connectome? At least in the constrained simulated examples we considered.

Using the available experimental data [67, 19, 33] we constructed a structural geometric connectivity network model (Methods 4.1) in order to analyze the dynamics on the C. elegans connectome using our dynamic signaling framework. This is in direct contrast to other attempts to model C. elegans using biophysical models [61, 57, 36, 38]. Biophysical models are computationally expensive [27] and require parameter estimates and assumptions that are not typically readily measurable or known in most cases. For example, ion-channel inactivation times and synaptic weights, for which there is little experimental data [44, 37]. In contrast, the dynamic simulations that underlie the analysis in our work here required only four considerations - an estimate of the signal conduction velocity, an estimation of a node refractory period, node type, that is, whether the node is either inhibitory or excitatory, and spatial node locations and connectivity, which are known [58]. Each node (i.e. neuron) in the network was assigned a 2 dimensional coordinate location, based on the known anatomical data, and we assumed edges were straight line connections between nodes.

We specifically studied the dynamics which resulted from stimulating a chemosensory neuron (ASEL) in a known feeding circuit. Experiments have shown that the activation of ASEL ultimately results in forward locomotion [60, 65], which is produced by the activations of mid-body motor neurons [32, 69]. First, we showed that the spatial embedding of the feeding circuit within the entire C. elegans connectome increases the dynamical repertoire of the circuit. We did this by quantifying the number of unique network states resulting from stimulation of ASEL in spatially-aware, i.e. spatially defined connectome, and spatially-unaware networks. Interestingly, the dynamics on the spatially-aware network that resulted from chemosensory activation, resulted in contralateral motor neuron activations in ventral (VB) and dorsal (DB) classes of motor neurons. This rhythmic alternating back and forth motor neuron firing pattern is qualitatively indicative of that required for movement of the worm. This result is subtle but critical in its interpretation. It is important to realize that we did not in any way model or intentionally emulate such rhythmic oscillatory dynamics between these two motor neuron populations. This dynamic behavior was in response to a single impulse input into the network (via a single activation of ASEL), and reflects the inherent (and we propose) purposeful structural wiring of the C. elegans connectome that has evolved over a very long time to serve purposeful behavioral functions.

Next, we analyzed and identified the causal neuronal signaling paths beginning at ASEL to VB and DB neurons. This reflects a unique part of the work and methods, enabled by the way our dynamic framework and simulation model were constructed. Neuronal signaling paths encode the causal chain of node activations over the course of the dynamics on the network. To describe each signaling path we introduced the notion of a Temporal Sequence (TSeq). Each TSeq is a temporally ordered sequence of nodes, formally a walk on a graph [22]. We determined the efficacy of sub-networks derived from the full network, through a quantitative comparison of TSeqs. More generally, TSeqs can be used to study the affects of network geometry (based on spatial embedding of network elements) and connectivity on dynamics. This is a key challenge in the analysis of complex geometric networks much more broadly than just an analysis of C. elegans. And as such, the methods we develop here are universally applicable to any spatial-temporal geometric network with signaling latencies. Furthermore, we decompose the complex network activity into a basis set of TSeqs which in turn form all the observed network activity. This basis set of TSeqs can be used as a signature of the dynamics on the network and they can also be used to construct sub-networks which more closely resemble the dynamics of the larger network. Finally, using TSeqs we show the interaction of classes of neurons from the motor neuron circuit influencing VB and DB activity.

More generally, the computational workflow we develop here can be used to generate experimentally testable hypothesis [23] regarding concurrent signaling paths supported by a connectome. TSeqs can also be used to generate a set of candidate sub-networks that preserve signaling paths, upon which more computationally expensive models can be implemented. Other on-going work in our group using these methods includes the analysis of causal dynamics on complex networks for machine learning. For example, in a general network setting where we wish to encode images through the causal dynamics, we can analyze the statistics of the resultant TSeqs, and other mathematical objects, such as, causal network motifs [47, 2, 68, 20]. Thus, these methods extend beyond computational neurobiology to potential applications to new forms of machine learning as well.

## 2 Results

### 2.1 The contribution of network geometry on dynamics

Geometric information plays an important role in network dynamics [58, 12]. Endowing edges with signaling delays that result from their physical geometry, i.e. convoluted paths in space, adds significant richness to the resultant network dynamics due to the offset in signal arrival times at nodes and the subsequent effects that has. Within the context of our model, we quantified the affect of signal delays on network dynamics in C. elegans. Signal delays are directly related to the spatial location of each of the neurons in the network. To do this comparison, we compared the network activity of the spatially embedded C. elegans connectome (Figure 1A), henceforth referred to as the Full network (Methods: 4.1), to the activity resulting from a latticed version of the C. elegans network, which we refer to as the Lattice network (Methods: 4.3). While the Lattice network’s edge connectivity and node types (inhibitory or excitatory) are identical to the Full network, we set the Lattice network’s edge signal delays to be a constant value; this then eliminates any affects of spatial embedding on the dynamics.

**Figure 1:**
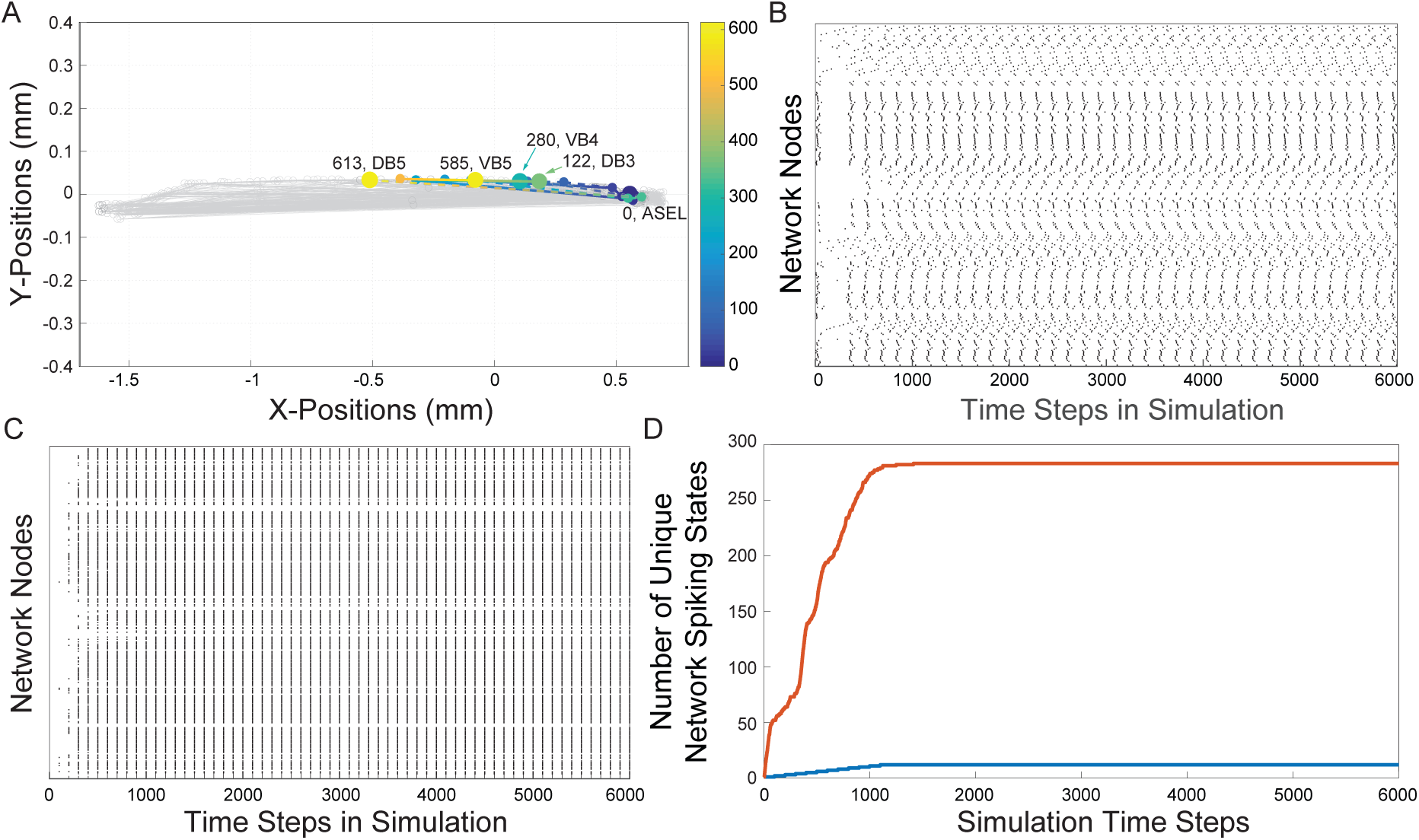
**(A)**, The reconstruction of the C. elegans connectome. Each gray line represents an edge. Each edge represents chemical connections between nodes. The colored dashed lines represents a sample of the traversal of a signal from the sensory neuron ASEL through the interneurons in the network to the VB and DB classes of motor-neurons. The color gradient indicates the time course as the signal traverses the network. Cold colors represent early in the neuronal signaling pathway and warm colors represent late in the neuronal signaling pathway. The node labels and numbers next to them represent the names of the motor neurons at the terminus of the sequence of activations and the time step when that terminal node was activated. **(B)**, The Full Network’s spike raster resulting from ASEL stimulation. The activity is a direct result of the signaling parameters (signal conduction velocity, refractory period), network connectivity, and the spatial locations of each of the nodes. Each dot represents the activation of a node in the network, the white space represents an inactive state of the node. Each discrete height on y-axis represents a specific node in the network. **(C)**, The Lattice Network’s spike raster caused by stimulating ASEL. The Lattice Network preserves the signaling parameters and network connectivity, but in effect modifies the spatial locations relative to the Full Network. We set the delays for all edges in the Lattice Network to be the same. The resulting spike raster is significantly less varied than that of the Full Network. We will quantify this previous statement with the next panel. **(D)**, Each curve is a cumulative count of the unique number of network states of presented by the Full (red curve) and Lattice (blue curve) Networks as they evolve over time. We observe an approximately a 20 times more network states in the dynamics of the Full Network than dynamics of the Lattice Network.

We compared the network activity between the Full and Lattice networks that resulted from a single pulse stimulus of the ASEL neuron (Figure 1B,C). This effectively represents the simplest stimulus and input into the network that is possible. In the Lattice network, we observed that the network activity went through a transient period of node activations, but which eventually results in almost all nodes firing at the same time in a periodic manner (Figure 1B). In contrast, although the Full Network’s node activity also went through transient and periodic activity phases, we observed a greater number and variability of patterns in its network state repertoire (Figure 1C). To quantify the variability in network activity that resulted from the geometry, we counted the number of unique network states. We describe the network’s state through a vector of nodal states at each discrete simulation time point. Concretely, if the state of node *i* at time *t* is given by *y*_*i*_(*t*) = {0, 1}, then the state of a network, with *N* nodes, at time *t*, can be written as **y**(*t*) = {0, 1}^*N*^. In Figure 1D we show the cumulative count of unique network states over the course of the simulation. The Lattice network assumed 12 unique states, while the C. elegans network assumed 280 unique states across the sampling points (we did not consider permutations of consecutive states). These results suggest that the geometry of a spatial-temporal network is an critical consideration to the dynamical range of the system. Throughout the rest of this paper we only make use of the spatial geometry of the C. elegans connectome, i.e the Full Network.

### 2.2 Qualitative comparison of network activity between the Full and Feed Networks using temporal sequences

The behavioral consequence of the activation of the ASEL neuron is the forward locomotion of the worm [69]. This is achieved by the resultant synchronized contralateral periodic firing of alternating populations of VB and DB motorneurons in the feeding circuit, which we will refer to as the Feed network throughout the rest of the paper. Its construction is detailed in the Methods: 4.2. Thus, we focused our analyses on the qualitative firing patterns of VB and DB populations in our simulations following single impulse stimulations of ASEL by mapping casual TSeqs. Additionally, because sub-networks are commonly used to study neighborhoods of complex networks, not only were we interested in the patterns of activity produced by the feeding circuit in isolation, identifying TSeqs from ASEL through interneurons to VB and DB motorneurons, but we also investigated whether sub-network patterns in the feeding circuit were preserved when we mapped TSeqs resulting from ASEL activation for the feeding circuit embedded in the complete C. elegans connectome, i.e. the Full Network.

We compared the network activity of the Feed and Full Networks to a set of respective null model networks (Methods: 4.5). Null model networks preserve the total number of nodes, node types, and node locations, but do not preserve the edge connectivity. The number of edges in the null model networks were approximately the same, but the edge connectivity was randomized. To do this, we computed TSeqs in order to identify the neuronal signaling paths which lead to motor neuron activations in the Feed, Full, and their randomized Null network versions.

After stimulating ASEL in the Feed and Full Networks, and waiting for the network activity to stabilize, we plotted a histogram of the resultant activity of VB and DB motor neuron classes over that time interval of the simulations. We observed that the VB and DB motor neurons activated in a staggered back and forth synchronized contralateral periodic manner (Figures 2A,C). This periodic activity pattern is qualitatively similar to the contralateral firing patterns necessary for the locomotion of the worm [71, 32, 53]. We observed that both the Full and the Feed network preserved the general patterns of ventral and dorsal neuronal activity.

**Figure 2:**
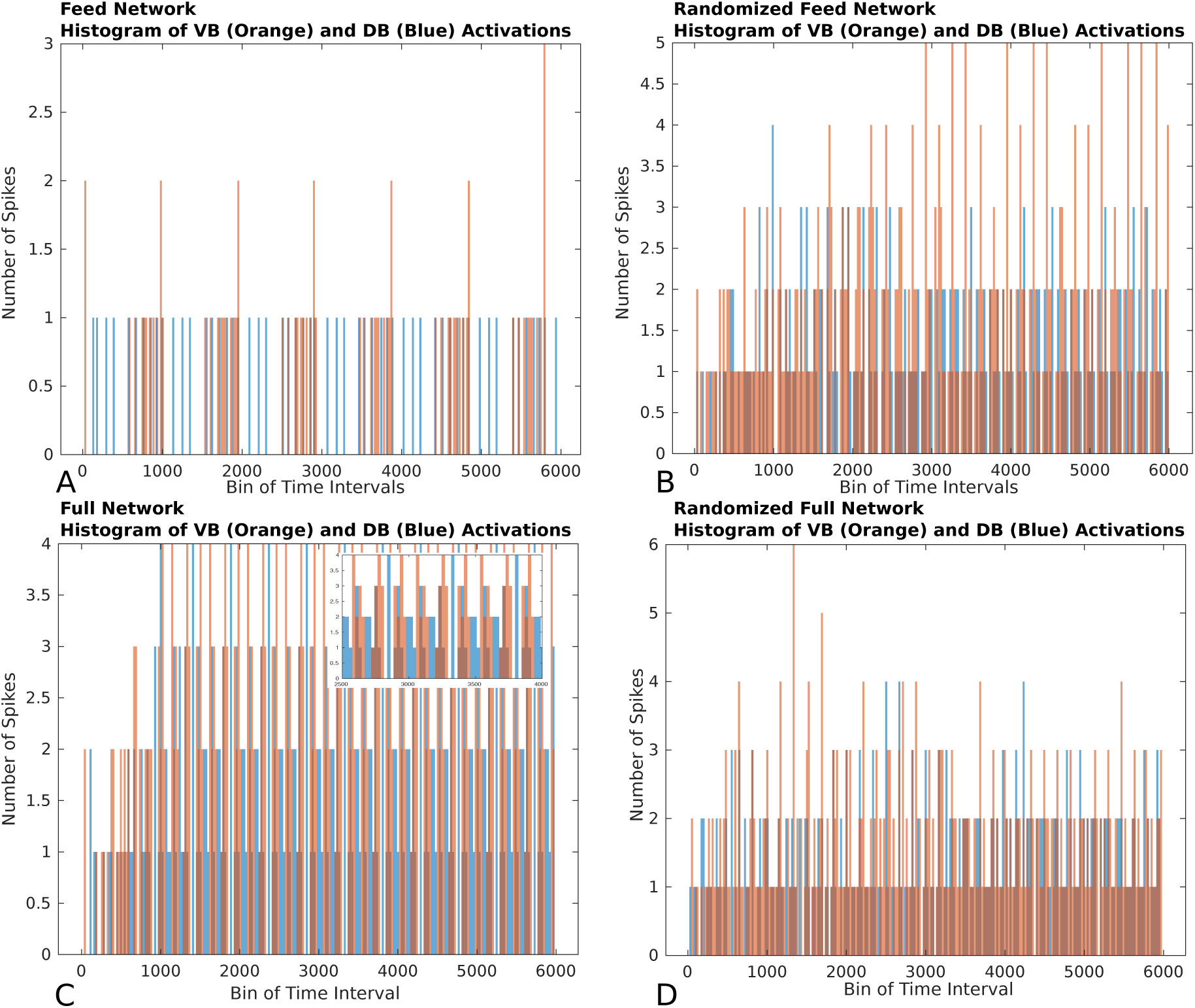
Histograms of the number of firings of nodes from the VB (orange rectangles) and DB (blue rectangles) classes. The bin size of the histogram is 250 time steps. **(A)** Histogram of the Feed Network’s VB and DB class motorneurons activations. We observe that the VB and DB classes of motorneurons activate in an alternating pattern. **(B)** Histogram of the Gilbert Randomized Feed Network’s VB and DB class neurons activations. Because the randomization procedure significantly modified the network connectivity, the class based alternating neuronal activation pattern was decimated. We observe a semblance of periodic node activations from the VB and DB classes after approximately 3000 time steps. **(C)** Histogram of the Full Network’s VB and DB class neurons activations. The C. elegans connectome gives rise to significantly more activity in the VB and DB classes of motor neurons in comparison to the activity in the Feed Network, Panel A. The dynamics on the Full Network also supported alternating VB, DB class activity. Because of the density of node activation, in the inset we show a zoomed in view of a sub-interval of activity to better view the VB-DB class alternating neuronal activation pattern. **(D)** Histogram of the Gilbert Randomized Full Network’s VB and DB class neurons activations. The randomization procedure resulted in the destruction of the class-wise alternating neuronal activation pattern.

Next, we studied what happened when we disrupted the edge connectivity in the Null networks. We used a randomization procedure [28, 24] that preserved the spatial locations of the nodes but replaced the original set of edges with a new set which was some subset of the total number of allowed edges. Despite the randomization, the connectivity density of the original networks were such that paths would eventually form and connect the ASEL sensory neuron to the VB and DB motorneuron populations for Null networks derived from both the Feed and Full Networks. Furthermore, the motorneuron populations always settled into periodic activity patterns. However, unlike the Feed and Full Networks, coordinated and synchronized contralateral alternating firing patterns were completely lost (Figures 2A,C). This empirically emphasizes the putative intentional design of the underlying wiring of the C. elegans connectome towards achieving an important behavioral function. The structural connectivity of the connectome is not random, but has likely evolved in a purposeful way to subserve specific purposes.

To understand how a single pulse stimulation of the ASEL neuron resulted in the firing patterns of in the VB and DB classes, we decomposed the network activity into TSeqs (Methods: 4.7). A TSeq represents the causal nodal interactions between a set of start nodes and a set of end nodes. Each TSeq is a walk, of an individual signal, on the graph from some start node to some end node. TSeqs capture the signaling pathways of the network, as such, they enable charting the course of a stimulus or any signal through the connectome. Only a subset of all possible walks on a graph are realizable at any given time. This is due to network parameters such as delays along edges, refractory periods of the nodes, and concurrent network activity. We focused on the TSeqs which start at the ASEL neuron, and either terminate or traverse the ventral, VB, or dorsal, DB, classes of motor neurons. We only computed TSeqs form ASEL to the VB and DB neuronal classes to ascertain relevant neuronal signaling paths (NSPs).

To better visualize TSeqs, we created a TSeq plot (Methods: 4.8), that showed the traversal of a set of TSeqs over time. On the TSeq plot the abscissa is time, and the ordinate is node number, similar to a raster plot. Each curve in the TSeq plots traces the causal walk of a signal on the structural connectome leading to the activation of a neuron of interest. The locations of the motor neurons (end nodes) on the TSeq plot are indicated by the dark blue horizontal lines. All the curves that are active in a time interval, trace the concurrent neuronal signaling pathways, which lead to subsequent activations of motor neurons from the neuronal classes of interest. To reduce the visual clutter in the TSeq plot we do not explicitly mark the nodes on the walk. The activated nodes are generally located at inflection points along the curve.

As anticipated based upon observing the histogram of node activity in the randomized network (Figures 2B,D), the TSeqs plots of the randomized networks show a complete breakdown of NSPs (Figures 3B,D) relative to TSeqs from the Full and Feed networks (Figures 3A,C). A notable qualitative difference between the TSeqs of the edge randomized and original networks was that the TSeqs of the Full and Feed networks enter a periodic regime of activity significantly sooner. Additionally, the TSeqs in the randomized network displayed convoluted neuronal signaling pathways relative to the orderliness of signaling pathways of the non-randomized networks.

**Figure 3:**
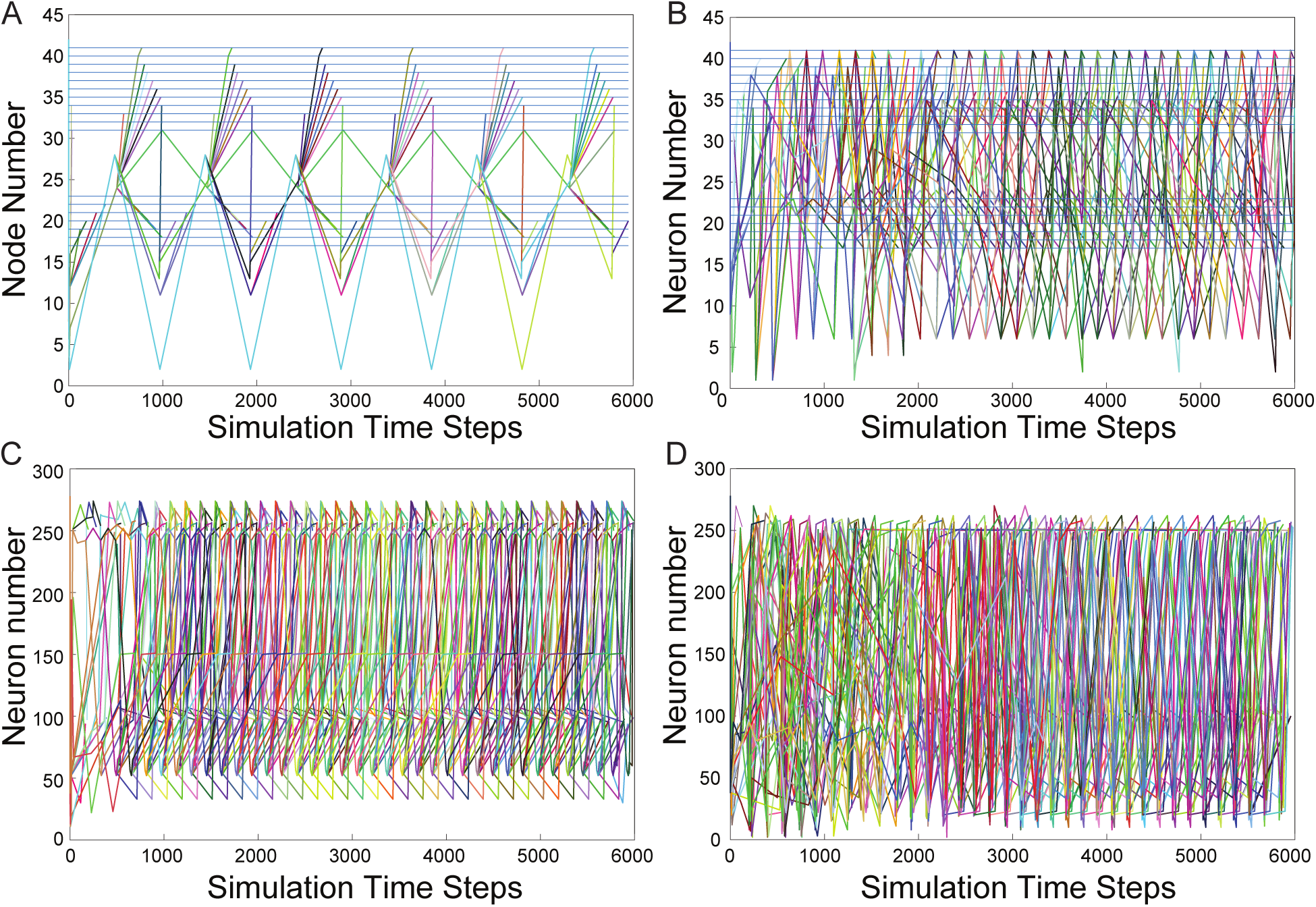
TS Plots. All TSs from ASEL to the VB and DB classes. **(A)**, TS plot of Feed Network. The activation patterns of neurons in the VB and DB classes are synchronized and staggered eventually giving rise to periodic sequences of node activity. **(B)**, TSs of the Gilbert Randomized Feed Network. Synchronized and staggered activations from the VB and DB class are absent. Transient period of network activity is longer than in Panel A. The TSs eventually enter a periodic sequences of node activity. There were significantly more TSs resulting from the randomized network in comparison to the TSs from the Feed Network as in Panel A. **(C)**, The number of TSs resulting for stimulating ASEL in the Full Network were significantly more than those from the Feed Network. Furthermore, the number of motor neurons were activated at a faster rate than they were in the Feed Network. This indicates that there are many other signaling pathways or more active signaling pathways from ASEL to the VB and DB classes of motor neurons which are either not captured or not as quickly activated by the Feed Network. The TSs of the Full Network seem to enter periodic activity within the first 1000 simulation time steps. **(D)**, TSs from Gilbert Randomized Full Network. The TS displayed a significantly longer transient period before the network activity entered a periodic regime. The transient period lasted approximately for the first 2000 steps of network activity.

In Figures 3A,C we show the TSeq plots of the Feed and Full Network’s respectively. Although the histograms of node activations showed a similar trend in class-wise VB and DB motor neuron activations (Figures 2A,C), the TSeqs plots show that the NSPs in the Full Network are markedly different (Figures 3A,C). We were motivated by this observation to develop a quantitative approach to compare TSeqs.

### 2.3 Quantitative comparison of temporal sequences

We developed a quantitative method called the Temporal Sequence-Similarity Measure (TSeq-SM). Before discussing the results, in the following section we first introduce the TSeq-SM algorithm. Next we describe a set of networks, all derived from the Full network, for which we computed TSeqs and then compared them using the TSeq-SM algorithm. In the last subsection we discuss the TSeq-SM results.

#### 2.3.1 Similarity Measure

We used the TSeq-SM to quantitatively determine the similarity between any two sets of TSeqs. Sets of TSeqs can be derived from the same network or from different networks. Of the two sets of TSeqs being compared by the TSeq-SM, one of the sets must be a user defined reference network. The TSeq-SM is calculated relative to the length of each TSeq from the reference network. We then count the number of TSeqs meeting some matching criteria *α*, where *α* is the percent of the length of one sequence from a set matching some sequence from another set. The TSeq-SM is similar to graph edit distances [56], but is applied to an ensemble of walks.

We outline the Similarity Measure in Algorithm (1). The inputs to the algorithms are (*X, Y, α*). The output of this algorithms is the TSeq-SM value. The TSeq-SM result can be stated in words as follows: Given network *X* as the reference, *n*_*X*_ TSeqs out of a total of *m*_*X*_ TSeqs from network *X*, found a *α* match to some subset of TSeqs of out a total of *m*_*Y*_ TSeqs from network *Y*. Upon completion of this procedure, we can ascertain the percent of TSeqs from finding a *c_α_* match by normalization: *c_α_/m_X_*.

Concretely, lets say we wish to calculate TSeq-SM between network *X* and network *Y*, where *X* is the reference. The sets *X* and *Y* respectively contain the TSeqs from networks *X* and *Y*. Let the *i^th^* TSeq of *X* be denoted *x*_*i*_, and the *j^th^* TSeq of *Y* be denoted *y*_*j*_, both *x*_*i*_ and *y*_*j*_ contain some number of node labels, each sequence of node labels representing a sequence of causal activations. Each comparison of the TSeq is relative to a threshold *α* which is a percentage of total elements in a particular *x*_*i*_ which in-order match the sequence *y*_*j*_ (Algorithm 1, getTSSimilarityMeasure). For each *x* ∈ *X* we evaluate (1) for every *y* ∈ *Y*:

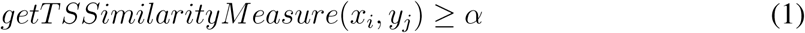

The output of (1) is either *True* or *False*. When (1) is *True*, a counter, *c_α_*, is incremented, and the next TSeq from *X*, *x*_*i*+1_ is checked against all *y* ∈ *Y*. The final value of the counter, *c_α_*, is number of TSeqs from *X* which met the criteria *α* relative to the TSeqs from *Y*. For example, if 4 out of 5 node activations in some *x*_*i*_ found an in-order match to some *y*_*j*_, and 4 out of 5 was the best match with respect to all *y* ∈ *Y*, then we say *x*_*i*_ meets the *α* = 0.8 criteria, and the counter *c*_0.8_ is incremented by 1.

#### 2.3.2 Networks Considered For Similarity Measure

We will list the networks we had used as input to the TSeq-SM algorithm and briefly describe how we generated each of them. There were 9 different networks in total. All networks that we considered were derived from the Full Network (Sec. 4.1).

As discussed in a prior section, the Feed Network contains a subset of the vertices of the Full Network. For the new set of vertices in the Feed Network, the edge connectivity their edge connectivity is preserved (4.2). In addition to the previously described Feed and Full Gilbert Randomized networks, we derived two more random networks using an Edge Swap randomization scheme (4.4). The two edge randomization procedures resulted in varying degrees of edge reorganization, while maintaining the number of nodes and node locations.

Of the two randomization procedures, the less drastic network reorganization is the Edge Swap based randomization because the network generation scheme prescribes that each existing edge swap their terminal node [22] according to some probability, therefore, not all edges need actually be different. The Edge Swap Random Network preserves various network parameters, such as, out-degree and network connectedness. Additionally, based on the number of iterations one chooses to run the Edge Swap procedure the network will steadily structurally deviate from the original. In contrast the Gilbert based randomization procedure’s only constraint is to maintain approximately the same number of edges between the original and the randomized networks. Therefore, the Gilbert based edge randomization [28] is a more drastic form of network reorganization.

The last set of networks we will discuss are called the Embedded Random Feed Networks (Methods: 4.6). These network preserves the edge connections of the Full Network outside of the Feed subnetwork. The subnetwork within the Full Network uses the same randomized edge connection profile as the standalone randomized Feed Networks. Two Embedded Random Feed Networks are generated, one based on the Gilbert Random Feed Network, the other based on the Edge Swap Random Feed Network.

#### 2.3.3 Similarity Measure Results

The results of the TSeq-SM are presented in Table 1. First, We will describe the organization of the Table of results, next the results themselves. The reference networks for each TSeq-SM are the antecedents in each of the column heading in the table. The top-half of Table 1 contains the TSeq-SM results of the Gilbert based randomization, and the bottom-half of the table contains the TSeq-SM results of the Edge Swap based randomization. In each half of the table, we show the TSeq-SM results resulting from each of the network pairs in the column heading. Stimulus was applied at the ASEL sensory neuron in all the networks, and the TSeqs of interest were all those which traversed the VB and DB sets of motor neurons.

**Table 1:**
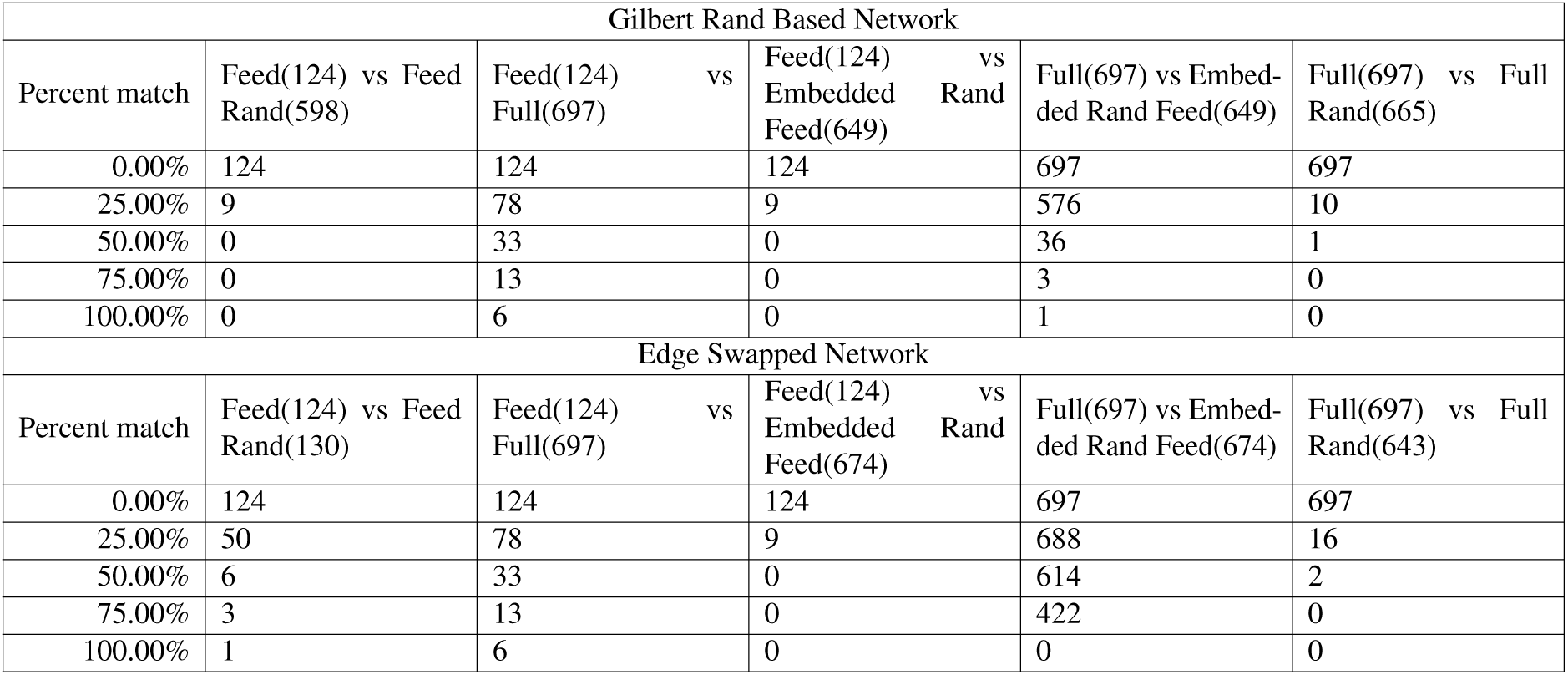
Table of Similarity Measures. The columns contain the pairs of networks compared. The rows in the leftmost column of the table represent the percent of a TS (the matching criteria *α* in Sec. 2.3.1) which matched any of the TSs from the reference network. The heading for each column are organized as reference network vs. target network. The cells in the table represent the number of TSs of the reference network which found a match TS from the target network meeting the corresponding row’s matching criteria.

Most generally, as the *α*-criteria increased, the number of TSeqs meeting the *α*-criteria decreased. This is expected as the variation in the structural network will cause variation in TSeqs. Therefore, fewer TSeqs from more structurally different networks will meet a higher *α* values.

The results indicate that as the structure of networks diverge the TSeq-SM values decrease. The TSeq-SM between the reference Feed Network and the Full Network was greater than the TSeq-SM between each of them and their edge randomized counterparts. This is because the Feed Network was more directly derived from the Full Network. Because of the less drastic network reorganization of the Edge Swap Networks, the TSeq-SM values of the reference Full/Feed Networks and their Edge Swap Randomized networks were higher than the TSeq-SM values of the Full/Feed Networks and their Gilbert Randomized counterparts.

Surprisingly, although the Feed Network was derived from the Full Network, the Feed Network in isolation only completely preserved 6 out of 124 TSeqs, i.e. *α* = 100%, the remaining 118 TSeqs followed different neuronal pathways. Ideally, the dynamics of a Feed Network should a subset of the dynamics of the Full Network. A natural question which arises is: Can we build a better subnetwork? Contained in the 697 TSeqs contain the necessary set of nodes required to construct a subnetwork. The sufficient set of nodes are those additional nodes involved in the neuronal signaling pathways of the concurrent network activity which ensures the existence of the 697 patterns. Determining the sufficient set of nodes is out of the scope of this paper because that requires iterative simulation and TSeq-SM measurement

Given the relatively poor TSeq-SM between the Feed and Full Networks, we were interested in quantifying the neuronal signal paths traversing nodes outside the Feed Network but in the the Full Network which link ASEL to the VB and DB classes of neurons. There are possible several approaches to generate an intermediate network which bridges the dynamical regime between the isolated Feed Network and the Full Network. One approach is to remove all the vertices and edges of the Feed Network from the Full Network, except those vertices associated with ASEL, VB, and DB. If we were to remove all the vertices of the Feed Network from the Full network certainly we would find TSeqs which lie completely outside the Feed Network. But removing all the vertices and edges associated with the Feed Network from the Full Network can drastically affect the overall dynamics of the network due to the absence concurrent dynamics. To workaround this issue, we used the Embedded Random Feed Network (4.6).

Relative to the TSeq-SM values resulting from the comparison of the isolated Feed Network (vs. Full Network), we observed higher TSeq-SM values while comparing the reference Full Network and the Embedded Random Feed Networks (Table 1, 5^*th*^ column). Although we found no TSeqs met the *α* = 100% criteria, across *α* values the Embedded Random Feed Networks better preserved TSeqs than the reference isolated Feed Network relative to the Full Network.

Although these results paint a cautionary tale for the analysis of isolated subnetworks, the quantitative approach of comparing network dynamics provides an indicator its efficacy. Further analysis of TSeqs provides an avenue for generating subnetworks, as well as preserving specific network interaction patterns. Coupling experimental observations and theoretical considerations with the TSeq-SM can provide a basis for choosing the constituents of a subnetwork.

**Algorithm 1.**
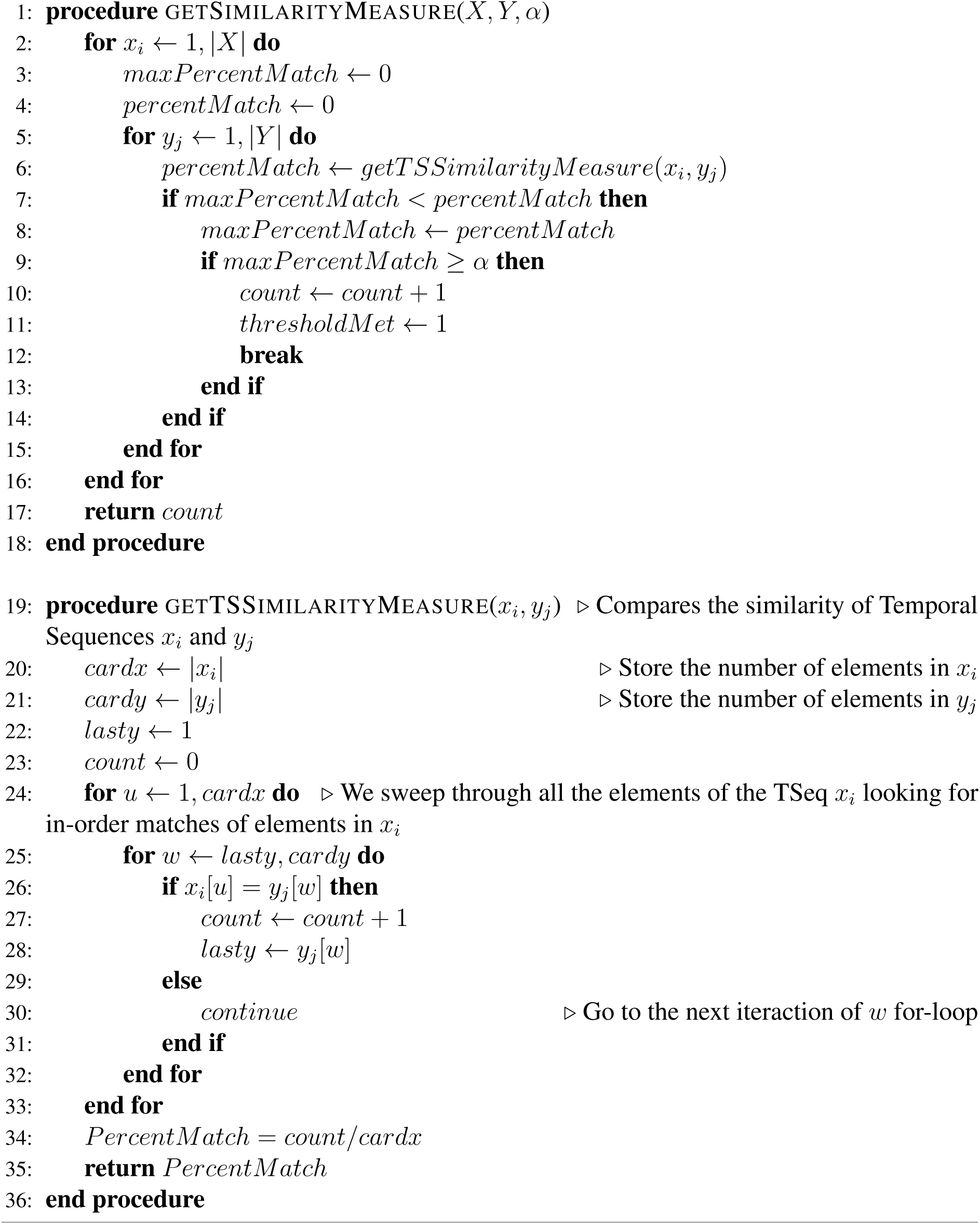
Similarity Measure Algorithm

### 2.4 Decomposing temporal Sequences into their basis sequences

We decomposed the complex patterns of network activity into a basis set of TSeqs (Methods 4.9). In brief, a basis set of TSeqs is a set of composed of TSeqs which arise from signals performing one-time walks and repeating walks on the graph. Repeated walk exist by virtue of signals traversing cycles of the graph. Each of the repeated TSeqs contained a set of sub-sequences, where each sub-sequence is repeated some number of times to compose other observed TSeqs. We can recompose all TSeqs from the set of One-Time walks and Repeated walks (given the repeating sub-sequence and their repetitions).

To ascertain the basis set of TSeqs, we categorized each TSeq into One-Time Temporal Sequences, and Repeated Temporal Sequences (Methods: 4.9). We found that the basis set of the 697 observed TSeqs from the Full Network contained 50 One-Time TSeqs, and 20 Repeated TSeqs. In Figures 4A and 4B we show the TSeq plot of One-Time and Repeated Sequences respectively. While the entirety of the One-Time Sequences are displayed in the figure, we only show the Repeated Sequences with a minimum of their repeating sub-sequences.

**Figure 4:**
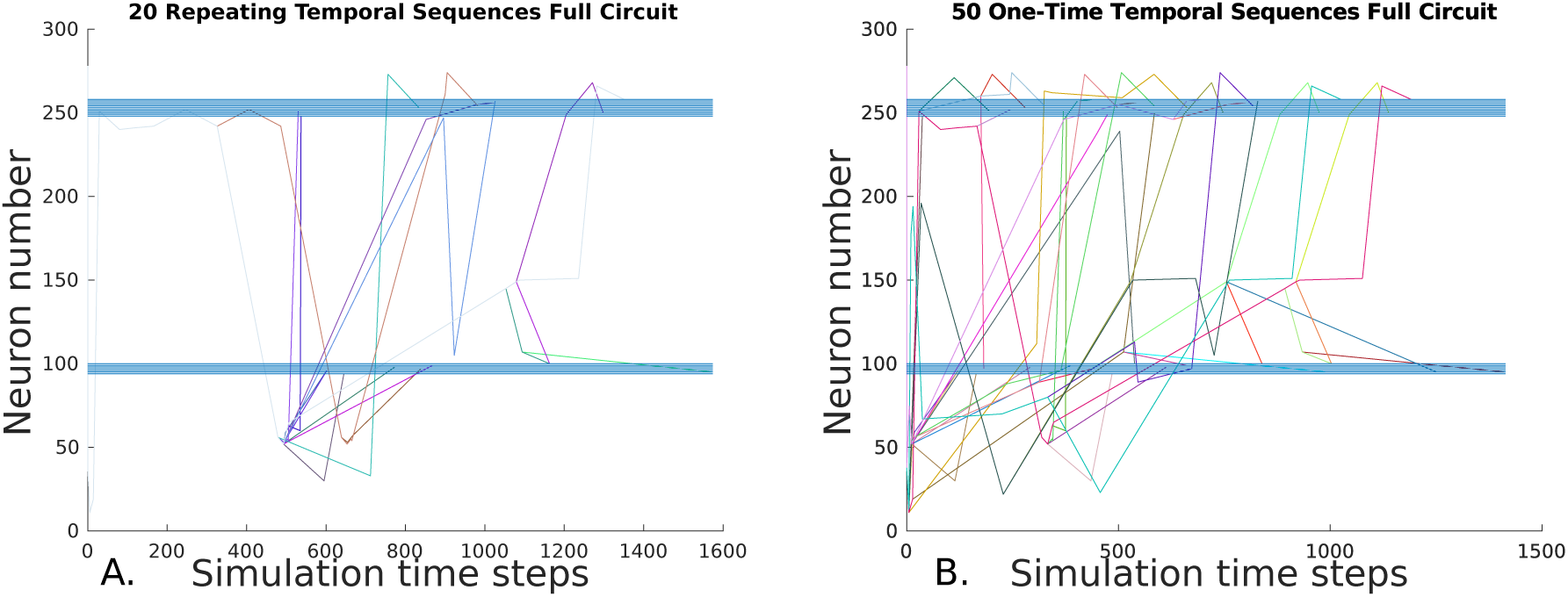
TS plots of the basis set of TSs from the Full Network. The basis set contains One-Time TSs and Repeated TSs. One-time TSs contain TSs which represent transient vertex activity on the network, and the Repeated TSs contain TSs which represent steady-state or repeating vertex activity on the network. **(A)**, TS plot of Repeated TSs. These TSs contain sub-sequences that when repeated some number of time is an exact match for another TS captured over the course of the network dynamics. **(B)**, TS plot of One-Time TSs. These TSs are unique neuronal signaling paths present in the dynamics.

Each of the 50 One-Time TSeqs are unique because they did not match any of remaining 696 TSeqs (not counting the TSeq being compared). Since there were 50 unique TSeqs, out of 697 total TSeqs, the 20 Repeated TSeqs compose the remaining 647 observed TSeqs through some sub-sequence repetition. Together the One-Time and Repeated Sequences form the minimal description of the dynamics and can be used as for further analysis and network construction.

### 2.5 Interacting motor neuronal classes along neuronal signaling pathways

Forward and backward movement of C. elegans results from the synchronized activations of banks of motor neurons [32] along its body. The cause of this synchronized activity involves the interaction of several classes of neurons [17, 71]. We analyzed TSeqs to identify the interaction between the various classes of mid-body motor neurons. We focused on TSeqs which started at the ASEL neuron, and traversed the mid-body motor neurons from the DA, DD, DB, VA, VD, and VB classes.

To determine the prevalence of various neuronal classes in neuronal signaling paths, we counted the number of TSeqs which contained various motor-neuron classes. As a starting point, in Figure 5A we show the number of TSeqs which contain neurons from individual classes. We found that all of the classes implicated in movement were present in at least some of the TSeqs (Figure 5A). Interestingly, the ventral side neuronal classes had significantly had more TSeqs traverse their neurons than the dorsal side neuronal classes.

**Figure 5:**
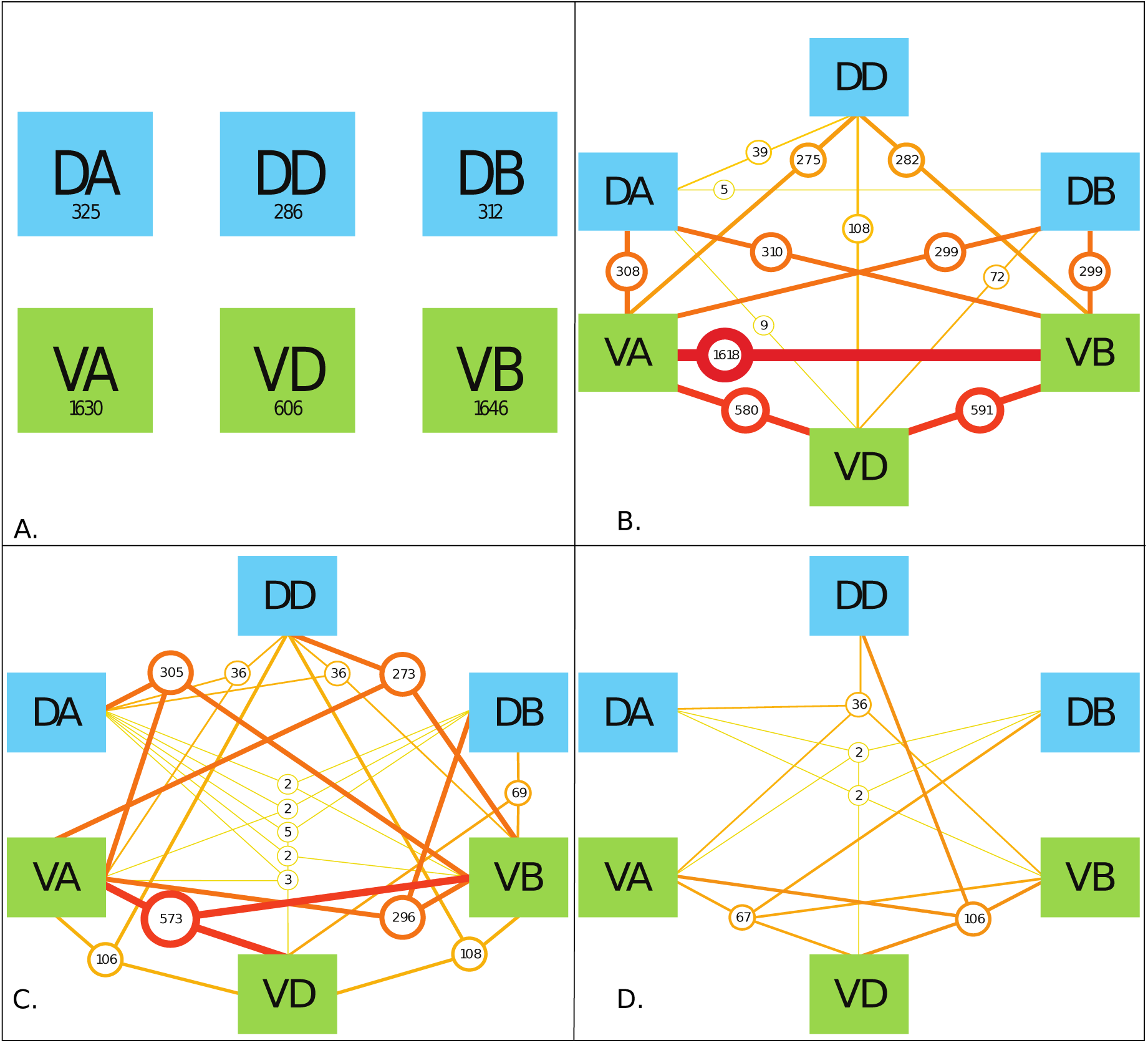
The interaction of various mid-body motorneuronal classes implicated in movement response upon stimulating ASEL in the context of the Full Network. We expand the motorneuron classes under consideration to include DA, DD, DB, VA, VD, and VB classes. We color code the dorsal side classes in blue and the ventral size classes in green. To elucidate the classes implicated in individual TSs, as a result of ASEL stimulation, we counted the number of TSs which contained at least one, two, three and four classes out of the six total classes under consideration. We only show 4 panels because there were no TSs which contained five or more classes in them. **(A)**, The values in each of boxes indicate the number of TSs with the specific neuronal class present within the TS. For example, looking at the box labeled DA, at least one of the neurons in the DA class was present in 325 TSs. The ventral classes were over-represented in TSs with single classes present. **(B)**, Here we considered the presence of any two classes of motor neurons in individual TSs. The most prevalent two classes present were neurons from the VA-VB classes. Only the DB-DD class interaction was not observed in any of the TSs captured. **(C)**, Here we considered three class interactions. The classes present in the TSs are given by the lines connecting the boxes. The number of sequences are given by the circles. Here we observe significantly fewer TSs than in Panel B. **(D)**, There are significantly fewer TSs meeting the four present classes criteria. We do not consider the order of present classes because of the much bigger combinatorial space to visualize.

Although the order of classes along the signaling pathways are relevant, they were not considered in this work because of the size of the permutation space. In Figures 5B,C,D we show the results for the number of TSeqs with 2, 3, and 4, neuronal classes present. Generally, as the number of classes present in a TSeq increased, the number of TSeqs decreased.

Our results on 3 and 4 class interactions, provides a future basis to experimentally test these neuronal interactions. Interestingly, individual TSeqs contained at most 4 interacting neuronal classes out of the possible 6 neuronal classes. Dynamics on the structural connectome limited the neuronal signaling paths to 4 classes of neurons and no more.

## 3 Discussion

A considerable body of work has approached the study of functional consequences from studying the structure of networks such as C. elegans [64, 62, 63, 13]. Particularly with C. elegans, detailed network analyses has focused on localized circuits [15, 29, 16, 71]. More recently, there a number of papers have looked at analyzing the dynamics at the whole brain scale to inform C. elegans research [35, 11, 70, 32, 1]. Our work here further contributes to this body.

We built a computational model of the dynamics of the C. elegans networks that uses our recently published framework which focuses on how local interactions among connected neurons give rise to the global dynamics in an emergent way to simulate the dynamics on C. elegans connectome [58]. We then computed TSeqs to analyze the underlying neuronal signal paths giving rise to the observed network activity. The analysis revealed that a single impulse stimulation simulating a food stimulus to the ASEL neuron recruits relevant populations of motor neurons with evoked neuronal activity patterns necessary for movement. Using TSs we charted the causal neuronal signaling pathways implicated in the response to relevant stimuli. Furthermore, our quantitative comparison of the network dynamics provided a quantitative method to design sub-networks. We proposed the Basis Sequences as the minimal description of the dynamics, which are not only compact signatures of the dynamics, but also provides guidelines for building networks that preserve dynamic patterns of node activity. The repeating set of basis sequences contain the pattern generators to sustain motor neuron activity. Finally, we quantified the extent and prevalence of various classes of mid-body motor neurons in individual neuronal signaling paths. These results suggest a number of interesting questions. For example, how do structural features and connectivity regulate the number of interacting populations? Why is there an imbalance between ventral and dorsal neuronal classes? And how well do the theorized class interactions align with experiments?

We intentionally focused on the simplest neuronal model that preserved key parameters significant in network signaling [58] in order to avoid getting bogged down by the biophysical complexity of neuronal networks. Future studies could extend our analyses of C. elegans in a number of ways. For example, by implementing a geometric framework with more complex neuronal models, using detailed biophysical signaling parameters, or making incremental modifications to the existing assumptions to uncover causal relationships supported by the network’s structure.

TSeqs based approaches can be implemented in different scientific domains where a network abstraction is possible and causal signaling dynamics can be discerned. Future work will investigate how the contributions of various factors affecting signaling dynamics, such as, edge/node models, network parameters and topology manifests itself in the causal relations between nodes in the networks to uncover how the network behaves at different temporal and spatial scales. Furthermore, we are interested in the stability and sensitivity of network dynamics to small perturbations and to variation in initial conditions.

Other work can extend the mathematical analysis of TSeqs. To gain insights into the structure of dynamics our approach for constructing TSeqs could be mapped to more complex mathematical objects such as a graphs [66, 4] and other topological structures [20]. Although we focused on the C. elegans connectome for this work, the analyses of TSeqs is agnostic to any specific network model.

## 4 Methods

### 4.1 Full Network Construction

Here, we will describe the computational model reconstruction of the C. elegans connectome [67]. We will call the model of the entire structural connectome the Full Network. All other networks in this work are derived from the Full Network. Mathematically, the network is a directed geometric graph [22, 58]. A graph *G* is a pair *G* = (*V, E*). Where *V* is the set of vertices/nodes/neurons, and *E* is the set of edges/link/axons such that *E* ⊆ *V* × *V*. A geometric graph is a graph whose vertices are embedded in euclidean space. A network is an applied instance of a graph, in our case a biological neuronal network. We consider directed networks because a neuronal action potential propagation is unidirectional, that is, a signal travel in one direction from one vertex to the next in one direction along an edge [34]. We used a simplified version of the Geometric Dynamic Perceptron model to describe the dynamics of each vertex [58]. In what follows we will discuss relevant network model parameters.

We constructed the directed adjacency matrix of the Full Network using publicly available connectivity information [67, 33, 3]. Let *A* be the network adjacency matrix, if there is a connection between neuron *i* and neuron *j*, then *A*_*ij*_ = 1, otherwise *A*_*ij*_ = 0. While neurons communicate to each other through multiple channels, our model only considered chemical connections between neurons, that is connections along axons. We did not consider electrical connections through gap junctions. Combining the structural connectivity with the location of neuronal cell bodies [19] we constructed the distance adjacency matrix. Out of the 302 neurons which make up the hermaphrodite C. elegans connectome, only 207 neuron locations are known [19]. Therefore, our final network consists of 277 vertices. We calculated the edge length using the euclidean distance between somatic bodies.

Although, there is much debate as to whether C. elegans fires action potentials, for simplicity, we assumed that some quanta of charge is transmitted between vertices through all-or-none stereotyped nodal events [45, 41, 40, 46], i.e. action potentials. Once a vertex initiates an action potential, the signal traverses all outgoing edges at a constant conduction velocity. We assumed a conduction velocity of 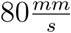 which is within the theoretical range for this type of organism [50]. We calculated the signaling time delays along edges through the geometrically derived edge lengths and the conduction velocity.

All vertices in our network network model are either inhibitory or excitatory. Excitatory vertices propagate excitatory signals through their edges. The response to an incoming excitatory signal is either the activation of the vertex receiving that signal, or the addition of potential to the receiving vertex’s membrane potential. If the receiving vertex’s membrane potential has exceeds threshold, it in turn propagates signals along its edges, and the vertex becomes refractory for the duration of its refractory period. The response to an incoming inhibitory signal is similar except no outing signals are generated by the receiving vertex. Only the GABA expressing neurons were considered inhibitory [26]. All other neurons types were considered excitatory [51], including the unknown neuron types. While a receiving vertex is refractory, no incoming signals will contribute its firing. The refractory period of all neurons are set to 4*ms* [8, 42].

Our network model uses a simplified version of the Geometric Dynamic Perceptron vertex model. Let the state of vertex *i* at time *t* be given by *y*_*i*_(*t*) = {0, 1}. When the vertex is in the active state *y*(*t*) = 1, when it is inactive *y*(*t*) = 0. While vertex *i* is initially in an inactive state it can instantly transition to the active state; but when the vertex *i* is in the active state it will become refractory for the time interval of the refractory period. We set the threshold for vertex activation to be one incoming signal. Therefore, when a signal arrives to the axon hillock of vertex *i* and it is in the inactive state, instantaneously, an outgoing signal will be generated and transmitted through all its outgoing edges.

We initialize network activity to quiescence. We initiate network activity by stimulating the ASEL neuron with a single pulse stimulus, eliciting outgoing signals based on the network connectivity. The simulation is run for approximately 1.5*s* of network activity, broken up into 6000 discrete time steps.

### 4.2 Feed Network Construction

The Feed Network [69] contains the neurons governing mid-body motorneuron response to feed stimulus at the ASEL chemosensory neurons. The Feed Network is subset of the Full Network. We reused the physical parameters from the Full Network for the Feed Network.

### 4.3 Lattice Network Construction

Our objective for constructing the Lattice Network was to study the affect of diminished influence of variation in edge lengths. We derived the Lattice Network from the Full Network We used the adjacency matrix of the Full Network in the Lattice Network. The difference between the Full and Lattice Networks is in the signaling parameters. We accomplished this by setting the edge signaling delays for all edges of the Lattice Network to 1*tu* (arbitrary time unit), and we set the vertex refractory periods to 0.9*tu*. This ratio of refractory period to signaling delay is considered to approach optimal from a local signaling perspective[58].

### 4.4 Edge-Swap Random Network Construction

We constructed a null model network called the Edge-Swap Random Network using the edge-swapping algorithm[54]. The starting point for the randomization procedure is the original network connectivity. The randomization procedure proceeds as follows: Two edges are randomly selected. If the two edges have two distinct source nodes and two distinct destination nodes, then the the two destination nodes are swapped, as long as the new edges from by the swap do not already exist. If any of the nodes associated with the randomly selected edges are the same, then two other edges are picked randomly, and the procedure is repeated. A network connectedness check is performed to make sure a separate network component is not created during the edge-swapping process. The number of times this process is repeated is approximately to the number of edges in the network. Each cycle through all the edges is consider an iteration of the algorithm. The code for the edge swapping algorithm can be found at [54].

We ran the algorithm for 10 iterations on both the Full Network and the Feed Network, to generate their randomized versions. This randomization scheme preserves the node degree distribution, by extension the number of edges in the network, and network connectedness. A new distance adjacency matrix is computed from the final adjacency matrix.

### 4.5 Gilbert Randomized Network Construction

Here we describe the randomization procedure we had used to generate the Gilbert Randomized Network. The primary difference between between the Full Network and their Gilbert randomized [28, 52, 24] counterparts is in the edge connectivity pattern and the new resulting edge lengths. The Gilbert random network was constructed based on the total number of edges in the Full Network. We calculated the probability of edge connection as follows: Let *N* be the number of nodes in the Full Network, and let *e* be the number of edges in the Full Network.

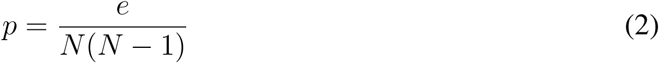

To create the random adjacency matrix, we create a matrix of random number between 0 and 1, then using *p* as the threshold value, we set the values of the matrix to 0 if the previous value was greater than *p* and 1 if the previous value was less than *p*. This results in an adjacency matrix of 0s and 1s with approximately the same number of edges in the Gilbert Randomized Network as in the Full Network. Using the positional data for the neurons, we calculated a distance adjacency matrix based on the resulting network connectivity. The signaling parameters of the Gilbert Randomized Networks were the same as the Full Network.

### 4.6 Embedded Random Feed Network Construction

We embedded the (Gilbert or Edge-Swap) edge randomized Feed Network into the Full Network to generate the Embedded Random Feed Network. We constructed this network by replacing the rows and columns of Full Network’s distance adjacency matrix with the values from the distance adjacency matrix of the edge randomized Feed Networks (see Sec. 4.4 and Sec. 4.5). All the signaling parameters of the Full Network were reused for the Edge Randomized Feed Network.

### 4.7 Temporal Sequence Construction

A Temporal Sequence (TS) is a specific type of walk on a graph [22]. The walk follows the trajectory of a quanta of signal between vertices along edges on a graph. In our network setting, TSs describe the trajectories of action potentials (discrete signals), along axons (edges), connecting neurons (vertices). The trajectory consists of a set of vertices traversed by the action potential emanating from the initial vertex activation.

Let us say we have a graph *G* = (*V, E*), which is a pair of sets formed by vertices, *V* and edges, *E*. Their elements are *v* ∈ *V* and *e* ∈ *V* × *V*. At time *t* = 0 lets say that node *v*_0_, some element in *V*, activates. Assume that *v*_0_ has an outgoing edge to *v*_1_, once the action potential from *v*_0_ reaches *v*_1_, *v*_1_ activates, which in turn generates an action potential. Lets say that this process continues until at some time *t* = *t*_*f*_ node 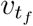 activates. The sequence of vertices 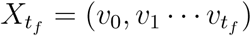 is the TS starting at *v*_0_(start node) traversing 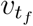 (end node). Generally, a set of TSs results between start node and end node in a time interval.

Since we were interested in TSs which traverse specific nodes in the network, we specified some start vertex (or set of vertices) for a TS and some end/traversed vertex (or set of vertices). In this work the set of start nodes contains only one element, ASEL, and the set of end nodes (more specifically traversed nodes) contain the neurons from the VB and DB classes. The resulting set of TSs represent the parallel paths through the graph traversed by signals beginning at the start node and traversing the end node over a time interval of interest.

### 4.8 Temporal Sequence Plot Description

We construct a plot similar to the raster plot to visualize the evolution of Temporal Sequences. We call this new plot a Temporal Sequence plot (TS plot). It has time steps on the x-axis, and node number on the y-axis. Each curve represents the causal trajectory of node activity, represented by a TS, over time. We separate each TS curve by color.

The activation of different parts of a network can result from shared neuronal pathways. Those TSs contain shared sub-sequences which are represented by overlapping curves on the TS plot. For example, in Fig. 4A we observe that all the Repeated TSs go through the same set of nodes at stimulus onset, before splitting into various Basis TSs.

In order to better visualize the paths through the nodes of interest, we drew blue horizontal lines on the TS plots. Each horizontal line is at the height of a neuron from to the VB and DB sets of neurons. In the Feed Network the DB neurons are located on the y-axis in the range from 17 to 23, while the VB neurons are located on the y-axis in the range from 31 to 41. In the Full Network the DB neurons are located on the y-axis in the range from 94 to 100, while VB neurons are located on the y-axis in the range from 248 to 258.

### 4.9 Basis Sequence Construction

The Basis Temporal Sequences (Basis TSs) are derived from the original set of TSs captured over the course of the dynamics of interest. Basis TSs consist of two sets of sequences, the set of One-Time Temporal Sequences (One-Time TSs) and the set of Repeating Temporal Sequences (Repeating TSs). First, we will give the intuition behind each of the constituent sets of sequences, then we will define them in more detail. Repeating sub-sequences result from signals traversing cycles on a graph. The vertices and edges involved in the closed walk can give rise to repeating network activity are candidate pattern generators of the network. Of the TSs captured over the course of the network dynamics, some result from graph cycles differ from one another only in the number of repeating sub-sequences. For each TS which differ in repeating sub-sequences, we create a compressed TS. We compose a set consisting of all the compressed TSs, and we name it the set of Repeating TSs. Given the description of the set of Repeating Temporal Sequences, we can readily describe the set of One-Time TSs as the remaining TSs which are not Repeating TSs.

Let *X* be the set of all TSs. Next, let *R* be the set of Repeating TSs in *X*, where *R* ⊆ *X*, and finally let *U* be the set of One-Time TSs, where *U* ⊆ *X*. We constrain the sets *R* and *U* such that *R* ∩ *U* = θ, that is, no TSs can be both a member of the Repeating TSs, as well as, the One-Time TSs.

Next, we will show how to ascertain whether some *x*_i_ ∈ *X* should be a member of *R*. The elements of *R* are such that for some *r*_i_ ∈ *R* and *r*_j_ ∈ *R*, the only difference between *r*_*i*_ and *r*_*j*_ is in the repetition of some sub-sequences between *r*_*i*_ and *r*_*j*_. Let us assume that *x*_*i*_ contains sub-sequences which repeat in different parts of its sequence. Let *I* = {*i*_1_*, i*_2_ *i*_*n*_} be the set of repeating sub-sequences of *x*_*i*_. Where each *i* ∈ *I* is the sub-sequence but without repeats, that is, each element of *I* is a primitive sequence. To account for the repeats, let *K* = {*k*_1_*, k*_2_ *k*_*n*_} be the set containing the number of repetitions of each of the primitive sequences in *I*. Now, any *x* ∈ *X* can be re-written as combination of primitive sequences together with its non-repeating sub-sequences. Concretely, given some TS *x*, we compress all its repeating sub-sequences in the form of 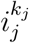, where *j* is the *j^th^* sub-sequence in *I*, which takes on *k*_j_ repetitions.

Every TS, *x_i_ ∈ X*, which can be derived from changing the number of repetitions of a primitive sequence, to match another sequence, *x*_j_ ∈ *X* is considered a Repeating TS. Both *x*_*i*_ and *x*_*j*_ are considered elements of *R*. Contained in the set *I* are the candidate pattern generators[30, 43] of the network, that is, the sequence of vertex activations which sustain ongoing network activity, and the realized cycles of the graph. It is not necessary that the dynamics realizes all the possible cycles of the graph. Because the elements of *R* are attained from observation, that Repeating TSs are not necessarily represent true periodic network activity, in the sense that it is guaranteed to repeat indefinitely.

Given the definition of Repeating Sequences, we can define the set *U* of One-Time TSs as *U* = {*u*|*u* ∈ *X*, *u* ∉ *R*}. The difference between *R* and *U* is that for all *x*_i_ ∈ *U* there is no other sequence *x*_j_ ∈ *X* such that the difference between *x*_*i*_ and *x*_*j*_ is only in some repetition of a sub-sequence in *x*_*i*_. Note *u*_i_ ∈ *U* may contain repeating sub-sequences. Since there are many approaches to ascertain Basis TSs, we do not provide an explicit algorithm to extract Basis TSs. It is possible to reconstruct all TSs in *X* form the the Basis TSs and knowledge of the number of repetitions of each sub-sequence.

